# *Vibrio cholerae*’s mysterious Seventh Pandemic island (VSP-II) encodes novel Zur-regulated zinc starvation genes involved in chemotaxis and autoaggregation

**DOI:** 10.1101/2021.03.09.434465

**Authors:** Shannon G. Murphy, Brianna A. Johnson, Camille M. Ledoux, Tobias Dörr

## Abstract

*Vibrio cholerae* is the causative agent of cholera, a notorious diarrheal disease that is typically transmitted via contaminated drinking water. The current pandemic agent, the El Tor biotype, has undergone several genetic changes that include horizontal acquisition of two genomic islands (VSP-I and VSP-II). VSP-I and -2 presence strongly correlates with pandemicity; however, the contribution of these islands to *V. cholerae*’s life cycle, particularly the 26-kb VSP-II, remains poorly understood. VSP-II-encoded genes are not expressed under standard laboratory conditions, suggesting that their induction requires an unknown signal from the host or environment. One signal that bacteria encounter under both host and environmental conditions is metal limitation. While studying *V. cholerae*’s zinc-starvation response *in vitro*, we noticed that a mutant constitutively expressing zinc-starvation genes (Δ*zur*) aggregates in nutrient-poor media. Using transposon mutagenesis, we found that flagellar motility, chemotaxis, and VSP-II encoded genes are required for aggregation. The VSP-II genes encode an AraC-like transcriptional activator (VerA) and a methyl-accepting chemotaxis protein (AerB). Using RNA-seq and *lacZ* transcriptional reporters, we show that VerA is a novel Zur target and activator of the nearby AerB chemoreceptor. AerB interfaces with the chemotaxis system to drive oxygen-dependent autoaggregation and energy taxis. Importantly, this work suggests a functional link between VSP-II, zinc-starved environments, and aerotaxis, yielding insights into the role of VSP-II in a metal-limited host or aquatic reservoir.

**Author Summary:** The Vibrio Seventh Pandemic island was horizontally acquired by El Tor pandemic strain, but its role in pathogenicity or environmental persistence is unknown. A major barrier to VSP-II study was the lack of stimuli favoring its expression. We show that zinc starvation induces expression of this island and describe a transcriptional network that activates a VSP-II encoded aerotaxis receptor. Importantly, aerotaxis may enable *V. cholerae* to locate more favorable microenvironments, possibly to colonize anoxic portions of the gut or environmental sediments.

## Introduction

The Gram-negative bacterium *Vibrio cholerae*, the causative agent of cholera (Harris et al. 2012), is well-adapted to two distinct lifestyles: as a colonizer of macroinvertebrates in the aquatic environment and as a potentially lethal pathogen inside the human intestine (Reidl and Klose 2002). *V. cholerae* persists in aquatic reservoirs by colonizing a variety of (mostly chitinous) biotic surfaces, such as copepods (Huq et al. 1983; de Magny et al. 2011; Tamplin et al. 1990; Sochard et al. 1979), shellfish (Twedt et al. 1981; Hood et al. 1981), and arthropods (Purdy and Watnick 2011; Broza and Halpern 2001). *V. cholerae* is ingested via contaminated drinking water or, less commonly, via undercooked seafood (DePAOLA 1981; Kaysner and Hill 2014). Once inside the human host, pathogenic varieties of *V. cholerae* (O1 and O139 serovars, (Chatterjee and Chaudhuri 2003)) rely on virulence factors to establish infection; the toxin co-regulated pilus (TCP) facilitates attachment to the intestinal wall (Thelin and Taylor 1996; Krebs and Taylor 2011) and cholera toxin (CTX) secretion ultimately drives efflux of water and salts from the intestinal epithelium (Sanchez and Holmgren 2008). CTX additionally promotes nutrient competition via depletion of free (i.e., not heme-bound) iron in the intestine (Rivera-Chávez and Mekalanos 2019).

The current (seventh) cholera pandemic agent, the O1 serovar El Tor biotype, arose from a non-pathogenic precursor via acquisition of TCP and CTX virulence factors (Hu et al. 2016). Unlike its pandemic predecessor (the Classical biotype), El Tor underwent several changes that include, among others (Son et al. 2011), the development of resistance against the antimicrobial peptide polymyxin B (Henderson et al. 2017; Hankins et al. 2012) and horizontal acquisition of two genomic islands (VSP-I and VSP-II) (Dziejman et al. 2002; O’Shea 2004). Presence of islands VSP-I and -II strongly correlates with pandemicity; however, only genes encoded on VSP-I have been directly linked to increased fitness in a host (Davies et al. 2012). VSP-II is a poorly understood 26-kb island that contains 24 ORFs (*vc0490-vc0516*) (O’Shea 2004), only two of which have validated functions: an integrase (*vc0516*, (Murphy and Boyd 2008)) and a peptidoglycan endopeptidase (*vc0503*, (Murphy et al. 2019)). The remaining uncharacterized genes are predicted to encode transcriptional regulators (VC0497, VC0513), ribonuclease H (VC0498), a type IV pilin (VC0502), a DNA repair protein (VC0510), methyl-accepting chemotaxis proteins (VC0512, VC0514), a cyclic di-GMP phosphodiesterase (VC0515), and 14 hypothetical proteins (O’Shea 2004). It is unclear if or how VSP-II enhances the pathogenicity or environmental fitness of El Tor. Intriguingly, VSP-II genes are not expressed under standard laboratory conditions (Mandlik et al. 2011), suggesting that their induction requires an unknown signal from the host or environment.

One signal that bacteria encounter under both host and environmental conditions is metal limitation. Bacteria must acquire divalent zinc cofactors from their surroundings to perform essential cellular processes; however, vertebrate hosts actively sequester zinc and other essential transition metals to limit bacterial growth (i.e. nutritional immunity) (Palmer and Skaar 2016; Kehl-Fie and Skaar 2010; Hood and Skaar 2012; Hennigar and McClung 2016). In the environment, *V. cholerae* frequently colonize the chitinous exoskeletons of aquatic and marine invertebrates and exposure to chitin oligomers has been suggested to induce zinc and iron starvation in *V. cholerae* (Meibom et al. 2004). In order to cope with zinc starvation stress, *V. cholerae* encodes a set of genes under the control of the well-conserved Zur repressor. When zinc availability is low, Zur dissociates from a conserved promoter sequence, allowing for expression of downstream genes. *V. cholerae* genes containing a known Zur binding region include those encoding zinc import systems (ZnuABC and ZrgABCDE) (Sheng et al. 2015), ribosomal proteins (RpmE2, RpmJ) (Novichkov et al. 2013; Panina et al. 2011), a GTP cyclohydrolase (RibA), and the VSP-II-encoded peptidoglycan endopeptidase (ShyB) (Murphy et al. 2019).

Here, we show that many genes of the VSP-II island are expressed during zinc starvation. These findings stemmed from an initial observation that a *V. cholerae* Δ*zur* mutant aggregates at the bottom of nutrient-poor liquid cultures. We hypothesized that this behavior was mediated by unidentified members of the Zur regulon. Using a transposon mutagenesis screen and RNA-seq, we identified Zur-regulated aggregation factors encoded on VSP-II. These included the transcriptional activator VerA encoded within the *vc0513-vc0515* operon. VerA (<u>V</u>ibrio <u>e</u>nergytaxis regulator A) activates expression of the AerB (<u>aer</u>otaxis B) chemotaxis receptor encoded by *vc0512*. We show that AerB mediates oxygen-dependent autoaggregation and energy taxis. Importantly, these results implicate a role for VSP-II encoded genes in chemotactic movement within zinc-starved environments.

## Results

### V. cholerae Δzur *mutant aggregates in minimal medium*

We noticed serendipitously that a *V. cholerae* N16961 Δ*zur* mutant (but not the wild type) aggregated at the bottom of a culture tube when grown shaking overnight in M9 minimal medium (**Fig. 1A; Movie S1**). *V. cholerae* aggregations in liquid culture are reportedly mediated by numerous mechanisms (e.g. quorum sensing, attachment pili (Jemielita et al. 2018; Taylor et al. 1987; Sun et al. 1997; Kirn et al. 2000; Adams et al. 2019; Trunk et al. 2018)) and stimuli (e.g. autoinducers, calcium ions (Jemielita et al. 2018), cationic polymers (Perez-Soto et al. 2018)), but none had been tied to zinc homeostasis. We therefore sought to identify factors that were required for Δ*zur* to aggregate. Aggregation (quantified as the ratio of optical densities (OD_600nm_) in the supernatant before and after vortexing) was alleviated by complementing *zur in trans*, excluding polar effects resulting from *zur* deletion (**Fig. 1B**). We next examined the role of zinc availability on aggregate formation. Since metals can absorb to the surface of borosilicate glass culture tubes (Struempler 2002), we instead grew *V. cholerae* in plastic tubes and noted that Δ*zur* still aggregated at the bottom (**Fig. S1A**), indicating that this phenotype is not linked to the properties of the culture vessel. Imposing zinc starvation via deletion of genes encoding *V. cholerae*’s primary zinc importer ZnuABC caused cells to aggregate similarly to the Δ*zur* mutant. Aggregation of Δ*znuABC* (which still elaborates the low-affinity zinc transporter ZrgABC (Sheng et al. 2015)) was reversed by zinc supplementation (**Fig. 1C**). In contrast, the Δ*zur* mutant, which constitutively expresses zinc starvation genes, aggregated in both the presence and absence of exogenous zinc. These data indicate that aggregation occurs in minimal medium when the Zur regulon is induced (i.e., during zinc deficiency or in a *zur* deletion strain) and is not a direct consequence of zinc availability *per se*. Surprisingly, none of the annotated members of the Zur regulon were required for aggregation (**Fig. 1D, Fig. S1B**), suggesting that there may be other Zur-regulated aggregation genes yet to be identified.

**Figure 1.**
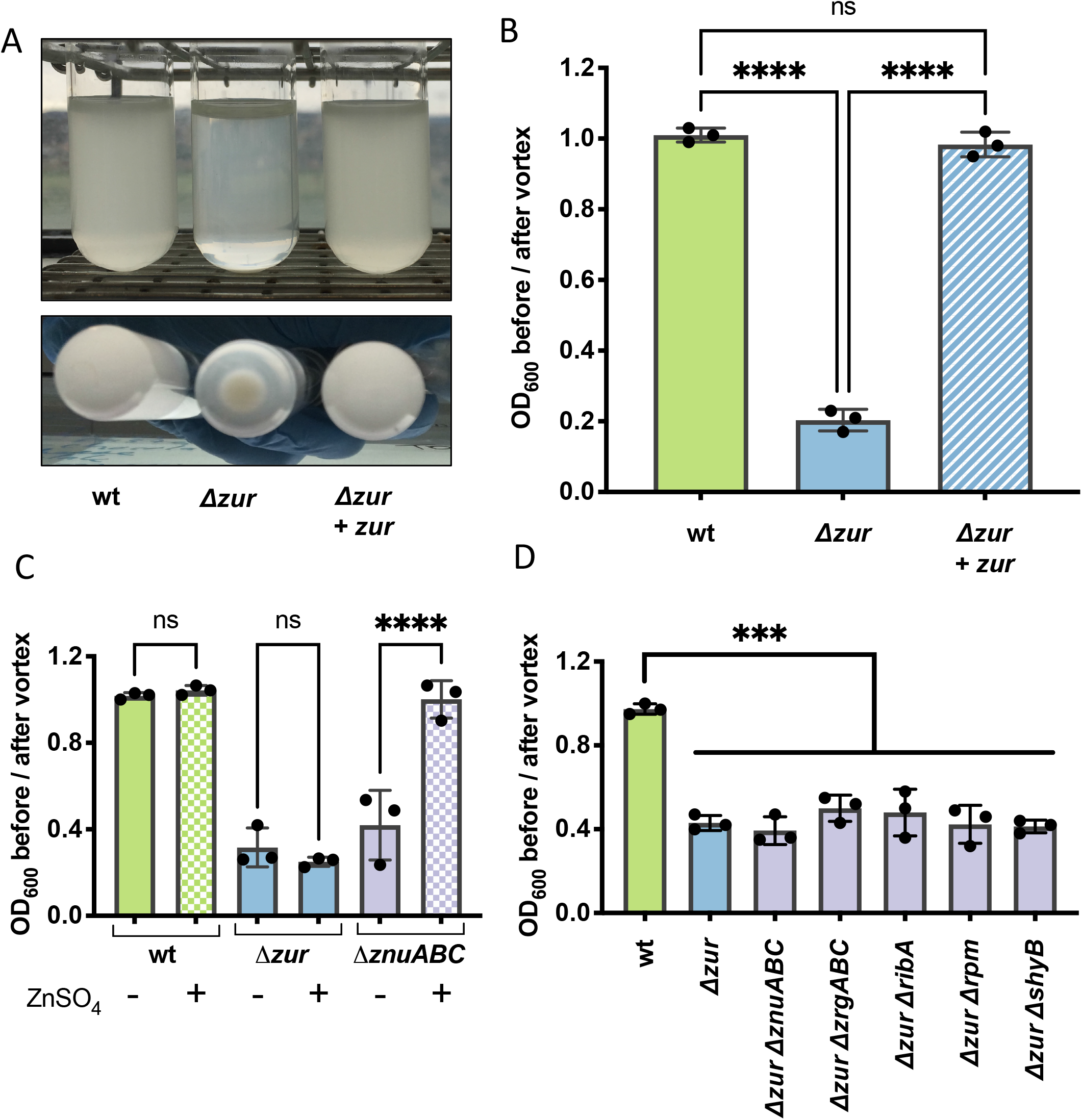
*V. cholerae* Δ*zur* mutant aggregates in M9 minimal medium. All strains were grown overnight in M9 minimal medium supplemented with glucose (0.2%). Aggregation was quantified by measuring the optical density (at 600 nm) of the culture supernatant before and after a brief vortex. A ratio close to 1 represents a homogenous culture, a ratio closer to 0 indicates aggregation (**A-B**) Representative images and aggregation measurements in medium containing inducer (IPTG, 500 µM) are shown for cultures of wild-type, Δ*zur*, and Δ*zur* carrying an integrated, IPTG-inducible copy of *zur* (denoted + *zur*). (**C**) Aggregation was measured in wild type, Δ*zur*, and a zinc importer mutant (Δ*znuABC)* grown in the absence (solid bars) or presence (checkered bars) of exogenous zinc (ZnSO4, 1 µM). **(D)** Aggregation was measured in wild type, Δ*zur*, and Δ*zur* lacking components of the zinc starvation response (*znuABC, zrgABC, ribA, rpmE2/rpmJ2*, or *shyB*). For all plots, the shown raw data points are biological replicates, error bars represent standard deviation, and asterisks denote statistical difference via Ordinary one-way ANOVA test (****, p < 0.0001; ***, p < 0.001; n.s., not significant).

### Δzur *aggregation requires motility, chemotaxis, and VSP-II encoded proteins*

We reasoned that we could leverage the Δ*zur* aggregation phenotype to identify novel components of *V. cholerae*’s zinc starvation response. To find such Zur-regulated “aggregation factors”, we subjected the Δ*zur* mutant to transposon mutagenesis and screened for insertions that prevent aggregation (see Methods for details) (**Fig. 2A**). Δ*zur* transposon libraries were inoculated into M9 minimal medium and repeatedly sub-cultured until no pellet formed. Transposon insertions sites were identified using arbitrary PCR on isolated colonies (O’Toole et al. 1999). The insertions that prevented aggregation overwhelmingly mapped to loci encoding motility and chemotaxis genes (**Fig. 2B)**. Twenty-four out of 48 recovered transposon mutants were disrupted in flagellar components or motility regulators. We reconstituted these types of mutations in Δ*zur* by inactivating flagellum assembly (major flagellin subunit, *fliC*) or rotation (motor protein, *motB*). Both Δ*zur* Δ*fliC* and Δ*zur* Δ*motB* failed to form a pellet (**Fig. 2C**), and aggregation could be restored by complementing each of these genes *in trans*. These data suggest that Δ*zur* aggregation is a motility-dependent process. Additionally, seven transposons inserted within genes encoding parts of *V. cholerae’s* chemotaxis machinery (*che-2*) (**Fig. 2B);** this system modulates bacterial movement in response to a chemical gradient. Mutating a component of this chemotactic phosphorelay (*cheA*::STOP) was sufficient to prevent aggregation in Δ*zur*, while *trans* expression of *cheA* restored pellet formation to the Δ*zur cheA::STOP* mutant (**Fig. 2C)**. Deletion of other *che-2* open reading frames also prevented Δ*zur* from aggregating (**Fig. S1C**). Collectively, these data suggest that motility and chemotaxis are required for Δ*zur* aggregation in minimal medium.

**Figure 2.**
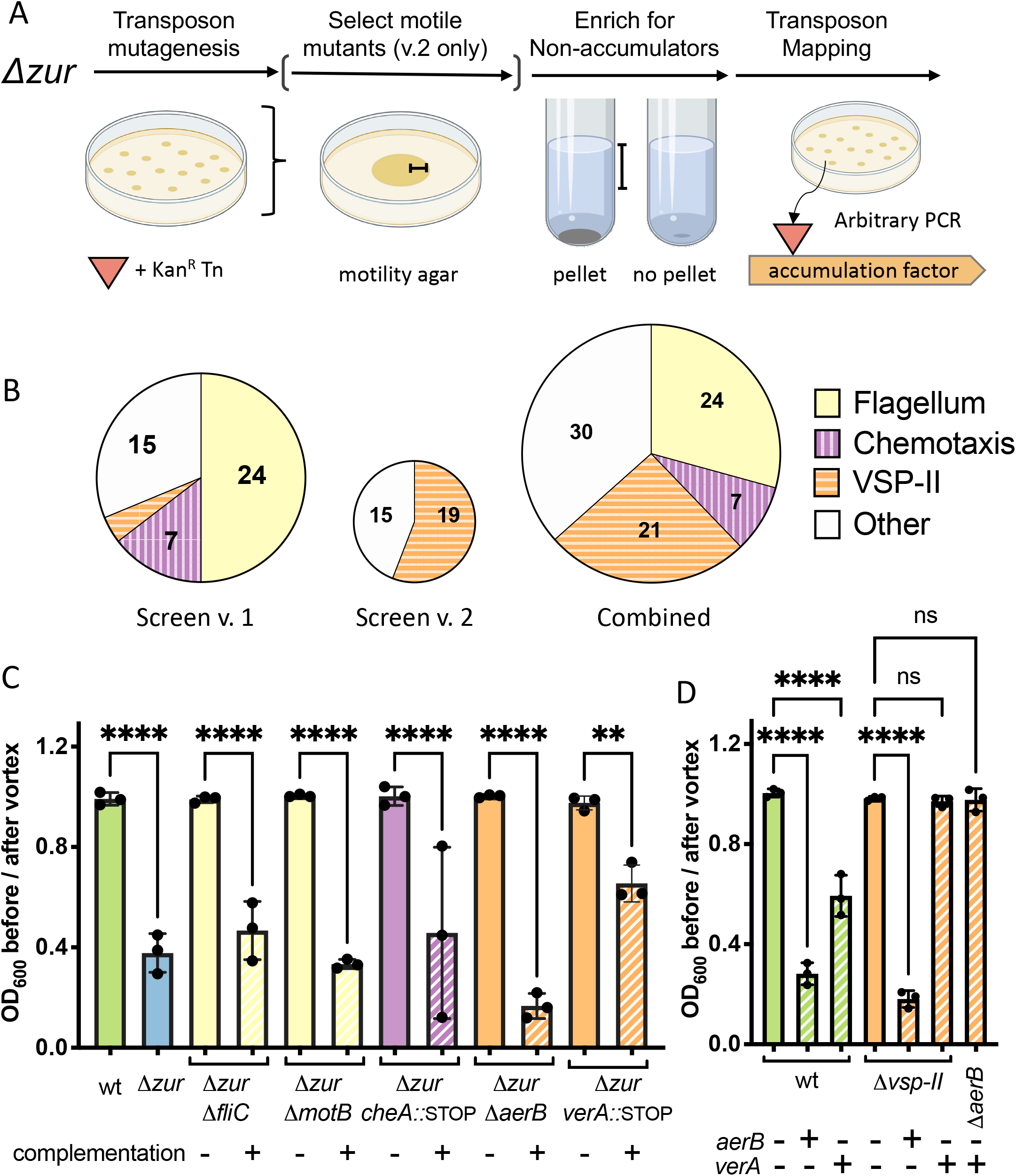
*zur* pellet formation requires motility and components of VSP-II. (**A**) Δ*zur* was mutagenized with mariner transposons to generate a library of insertion mutants (see Methods for details). Non-aggregating mutants within the library were enriched via repeated subculturing of the supernatant until no pellet formed. Transposon insertions were mapped using arbitrary PCR. In a modified version of this screen (v.2), the transposon library was pre-filtered to select for motile mutants on soft agar (0.3%). **(B)** Transposon insertions mapped to VSP-II genes (24 hits, orange/horizontal lines), genes encoding flagellar components and regulators (21 hits, yellow) and chemotaxis proteins (7 hits, purple/vertical lines). (**C**) Select motility (*fliC, motB*), chemotaxis (*cheA*), and VSP-II genes (*vc0512/aerB, vc0513/verA*) were mutated in a Δ*zur* background (solid bars) and complemented back *in trans* (+) under an IPTG inducible promoter integrated within chromosomal *lacZ*. (Note: P_iptg_-*verA* on a multicopy plasmid was used for complementing the Δ*zur verA::STOP* mutant. These cultures were grown with kanamycin to ensure retention of the control or *verA*-expressing plasmid). Aggregation in M9 glucose (0.2%) supplemented with inducer (IPTG, 100 µM) was quantified by measuring the optical density (at 600 nm) of the culture supernatant before and after a brief vortex. (**D**) Using the same approach described in **C**, chromosomal copies of *aerB* or *verA* were overexpressed in wild-type and Δ*vsp-II* (or a Δ*aerB*) background. Ten and 200 µM of IPTG were used for *aerB* and *verA* induction, respectively. For all plots, the shown raw data points are biological replicates, error bars represent standard deviation, and asterisks denote statistical difference via Ordinary one-way ANOVA test (****, p < 0.0001; **, p < 0.01; n.s., not significant).

We noted that the Δ*zur* phenotype resembles aggregation in *E. coli* rough mutants, which have reduced expression of lipopolysaccharides (Nakao et al. 2012). We observed similar aggregation in *V. cholerae* rough mutants (*vc2205::kan*), but this aggregation did not require motility to form (**Fig. S1D**) and is therefore mediated by a distinct mechanism. We anticipated initially that Δ*zur* pellet formation was a group behavior that may require processes associated with surface attachment (e.g., biofilm formation, attachment pili) or cellular communication (e.g., quorum sensing); however, such mutants were not identified by the transposon screen. We thus separately assessed this in a targeted fashion by testing whether Δ*zur* aggregates when deficient in biofilm formation (Δ*vspL*) or type IV pili attachment (Δ4: Δ*tcpA* Δ*mshA* Δ*pilA*, and orphan pilin Δ*vc0504*). Consistent with these processes not answering our screen, biofilm and type IV pili encoding genes were not required for Δ*zur* to aggregate (**Fig. SF**). Notably, N16961 contains an authentic frameshift mutation in the quorum sensing gene *hapR* (Heidelberg et al. 2000; Joelsson et al. 2006); however, a repair to *hapR* (Stutzmann et al. 2016) did not alter aggregation dynamics in Δ*zur* (**Fig. S1E**). We additionally demonstrated that other quorum sensing genes (Δ*csqA*, Δ*csqS*, Δ*tdh*, Δ*luxQ, or* Δ*luxS*) were dispensable for Δ*zur* aggregation (**Fig. S1F**). Taken together, these data indicate that Δ*zur* pellet formation is not a clumping phenomenon driven by typical colonization and aggregation factors, but rather a chemotaxis/motility-mediated assembly in the lower strata of a growth medium column.

Since aggregation appeared to require induction of the Zur regulon, we were surprised that the transposon screen was not strongly answered by genes with an obvious Zur binding site in their promoters. We reasoned, however, that our screen did not reach saturation due to the large number of motility genes encoded in the *V. cholerae* genome. We therefore refined the screen by pre-selecting for mutants that retained motility on soft agar, followed by a subsequent screen for loss of pellet formation in the motile subset of the mutant pool, as described above. Interestingly, 19 of the 34 transposon insertions answering this screen mapped to the Vibrio Seventh Pandemic island (VSP-II) (**Fig. 2B, Fig. S2)**, a horizontally acquired genomic region that is strongly associated with the El Tor biotype and the current (seventh) cholera pandemic. Transposons concentrated in a section of VSP-II that encodes a putative AraC-like transcriptional activator (VC0513, henceforth “VerA”), two ligand-sensing chemotaxis proteins (VC0512, formerly Aer-1 is henceforth referred to as “AerB”, and VC0514), and a cyclic di-GMP phosphodiesterase (VC0515). Notably, the *vc0513-vc0515* operon is preceded by a canonical Zur binding site and is thus a novel candidate for Zur-dependent regulation.

To validate the VSP-II genes’ involvement in Δ*zur* aggregation, we inactivated each gene in a Δ*zur* background through either clean deletion or, to avoid polar effects on other genes in the same operon, through insertion of a premature stop codon. Δ*aerB* and *verA*::STOP mutations prevented Δ*zur* from aggregating, whereas respective complementation with *aerB* and *verA* restored aggregation (**Fig. 2C**). To determine if these VSP-II genes were sufficient to generate aggregation, we overexpressed them in a wild-type and a Δ*vsp-II* background. Both *aerB and verA* overexpression caused the wild-type to aggregate, but only the *aerB* chemoreceptor triggered aggregation in a strain lacking other VSP-II genes (**Fig. 2D**). These data indicated that AerB drives aggregation and raised the possibility that VerA functions as a transcriptional activator of *aerB*. We additionally tested a deletion of the entire VSP-I island and mutations in all other open-reading frames on VSP-II (including *vc0514* and *vc0515*), none of which were required for Δ*zur* aggregation under the conditions tested (**Fig. S1G**). These two screens indicate that pellet formation in Δ*zur* is driven by chemotactic flagellar movement, with assistance from a VSP-II encoded transcriptional activator (VerA) and chemoreceptor (AerB). These results were intriguing given that very little is known about the regulation or function of VSP-II encoded genes.

### Several VSP-II genes are significantly upregulated in a Δzur *mutant*

Prior inquiry into VSP-II function was made difficult by a lack of native gene expression under laboratory conditions; thus, we prioritized mapping the transcriptional networks embedded in this island. Our data implies that the VSP-II genes of interest are expressed in Δ*zur*. Indeed, the *vc0513-vc0515* promoter region contains a highly conserved Zur-binding sequence approximately 200 bp upstream of the mapped transcription start site (determined by 5’-RACE in a Δ*zur* mutant, **Fig. 3A, S4A**). Although the distance between the Zur box and transcriptional start site is greater than that observed for most *V. cholerae* Zur targets, equivalent or greater distances are noted for the Zur-regulated *ribA* (140-210 bp upstream of the ORF) and *zbp* (∼380 bp upstream of ORF) in closely related *Vibrio spp*., respectively (Novichkov et al. 2013). To verify regulation by Zur, we measured *verA* promoter activity via a *lacZ* transcriptional fusion (*P*_*verA*_*-lacZ*), which encodes β-galactosidase (LacZ) and gives a colorimetric readout in the presence of a cleavable substrate (e.g., ONPG). As expected, transcription from the *verA* promoter in zinc-rich LB medium was robust in Δ*zur* relative to wild-type or a *zur* complemented strain (**Fig. 3B**). This data indicates that Zur negatively regulates *verA* transcription in a rich medium. We also tested *P*_*verA*_*-lacZ* expression in M9 minimal medium in a wild-type, Δ*zur*, and Δ*znuABC* background and noted that promoter activity corresponded to the conditions in **Fig. 1C** that triggered aggregation (**Fig. 3C**). For example, relative to a wild-type background, P_verA_ activity was robust in both Δ*zur* and a mutant deficient in zinc uptake (Δ*znuABC*). Consistent with Zur’s zinc sensing function, Δ*zur P*_*verA*_*-lacZ* strain retained high levels of β-galactosidase activity regardless of zinc availability, whereas *P*_*verA*_*-lacZ* in Δ*znuABC* was repressible with exogenous zinc. These data, in conjunction with the highly conserved Zur binding site, suggest that the VerA-encoding *vc0513-vc0515* operon is a novel component of the Zur-regulated zinc starvation response in N16961.

**Figure 3.**
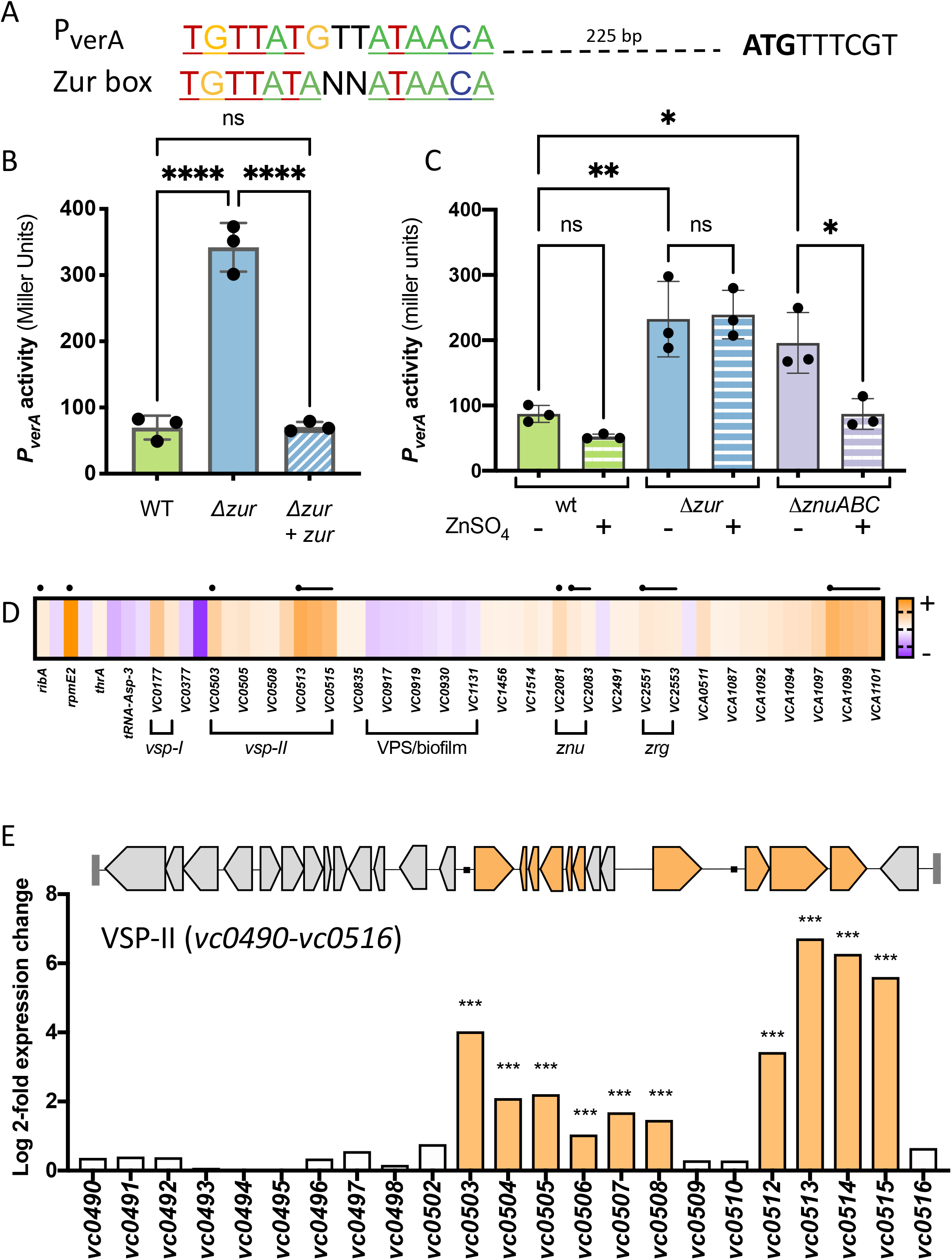
Several VPS-II genes are upregulated in a Δ*zur* mutant. **(A)** A predicted Zur-binding site (<u>TGTTAT</u>GTT<u>ATAACA</u>) located approximately 200 bp upstream of the *verA* open reading frame was aligned with the *Vibrionacae* Zur binding consensus sequence (Novichkov et al. 2013). Predicted start codon is indicated by bold “ATG”. **(B**) *P*_*verA*_*-lacZ* transcriptional reporters were introduced into a wild-type and Δ*zur* background, paired with either an empty vector or IPTG-inducible copy of *zur* (*+ zur*). Strains were grown overnight and diluted 1:100 into LB with kanamycin and inducer (IPTG, 200 µM). After 3 hours of growth at 37°C (to mid/late exponential phase), promoter activity was quantified in Miller Units by measuring β-galactosidase (LacZ) activity against an ONPG chromogenic substrate (see Methods for more details). (**C**) Wild-type, Δ*zur*, and Δ*znuABC* mutants carrying the *P*_*verA*_*-lacZ* reporter were grown in M9 minimal medium in the presence (+) and absence (-) of exogenous zinc (ZnSO_4_, 1 µM). After overnight growth (16 h), promoter activity was measured in Miller units. For bar graphs, raw data points represent biological replicates, error bars represent standard deviation, and asterisks denote statistical difference via Ordinary one-way ANOVA (****, p < 0.0001; ***, p < 0.001; **, p < 0.01; *, p < 0.05; and n.s., not significant). (**D-E**) RNA was isolated from cells at mid-log phase and prepared for RNA-seq (see Methods and Materials). Genes with significant differential expression in Δ*zur* (log 2-fold change > 1, adjusted p-value <0.05) relative to wild-type N16961 are shown. (**D**) The heat map indicates increased (orange) or decreased (purple) expression relative to the wild-type strain. Black circles represent putative Zur binding sites and lines represent corresponding operons. (**E**) Log 2-fold expression changes for all VSP-II genes (*vc0490-vc0516*) are shown alongside a schematic of VSP-II open reading frames. Black circles indicate present of canonical Zur binding sites.

Global transcriptomic studies of the Zur regulon have been conducted in a number of bacteria, but none thus far have been reported in the *Vibrio* genera (Hensley et al. 2012; Sigdel et al. 2006; Kallifidas et al. 2010; Gaballa et al. 2002; Bütof et al. 2017; Pawlik et al. 2012; Neupane et al. 2017; Mazzon et al. 2014; Moreau et al. 2018; Mortensen et al. 2014; Li et al. 2009; Lim et al. 2013; Pederick et al. 2015; Mastropasqua et al. 2017; Latorre et al. 2015; Schröder et al. 2010; Eckelt et al. 2014; Owen et al. 2007). We thus performed an RNA-seq experiment comparing transcript abundance in wild-type N16961 and Δ*zur* to assess *V. cholerae’*s Zur regulon more comprehensively (including indirect effects). To ensure sufficient repression of Zur targets in the wild-type, cells were grown to mid-log phase in LB medium. Analyses identified 58 differentially expressed genes in Δ*zur* (log2 fold change >1, adjusted p-value <0.05) (**Fig. 3D, Table S2**). Seven promoters (situated in proximity to 23 of the 42 upregulated genes) contained an upstream canonical Zur-binding site. Among them were known or inferred (based on *E. coli*) Zur regulon components, including genes that encode zinc uptake systems (ZnuABC, ZrgACD), an alternative ribosomal protein (RpmE2), and a GTP cyclohydrolase (RibA). This transcriptomic analysis also uncovered what appears to be a bidirectional promoter with a Zur box: this locus encodes a strongly upregulated ABC-type transporter (*vca1098-vca1101*) in one direction and upregulated portions of the chemotaxis-3 (*che-3*) cluster (*vca1091-vca1095, vca1097*) in the other. Using a *lacZ* transcriptional reporter, we verified that the ABC-type transporter is indeed Zur-regulated (**Fig. S3**). Neither the transporter nor the *che-3* cluster, however, were required for Δ*zur* aggregation (**Fig. S1F)**. We observed a striking cluster of nine up-regulated genes on VSP-II (comprising 35% of the annotated open reading frames on VSP-II), including the previously characterized peptidoglycan hydrolase ShyB (encoded by *vc0503*) and the *vc0513-vc0515* operon, consistent with our transcriptional fusion data and the transposon screen (**Fig. 3D-E**).

The RNA-seq analysis also identified 36 differentially expressed genes that lacked canonical Zur binding sites. Nineteen of these genes were significantly up-regulated in Δ*zur*, including several genes on VSP-II (*vc0504-vc0508* and *vc0512*) (**Fig. 3D-E, Table S2)** and VSP-I (*vspR, capV*). Other upregulated transcripts in Δ*zur* encode for cholera toxin (*ctxA/B*), the toxin co-regulated pilus biosynthesis proteins (tcpT/H), and a chitin binding protein (*gbpA*). Seventeen genes were significantly down-regulated in Δ*zur*, many of which were related to vibrio polysaccharide (VPS) synthesis and biofilm formation (Fong et al. 2010) (**Fig. 3D, Table S2**). Thus, a *zur* deletion affects numerous genes indirectly, possibly through Zur-dependent secondary regulators (e.g. VC0515, cyclic di-GMP phosphodiesterase; VerA, AraC-like transcriptional regulator), via secondary responses to the influx of zinc that the Δ*zur* mutant is expected to experience, or via Zur-dependent small RNA interference.

### *VerA is a Zur-regulated transcriptional activator of* aerB

Our mutational analyses above raised the possibility that the putative MCP AerB is controlled by the transcriptional activator VerA. Given the importance of AraC-family regulators in governing *V. cholerae*’s host-associated behaviors (e.g., ToxT, intestinal colonization and virulence (DiRita 1992; DiRita et al. 1991; Higgins et al. 1992; Weber and Klose 2011); Tfos, chitin-induced natural competence (Metzger and Blokesch 2016; Yamamoto et al. 2014; Dalia et al. 2014)) we sought to characterize the full VerA regulon. We performed an RNA-seq experiment comparing transcript abundance in N16961 overexpressing *verA*, relative to an empty vector control. Surprisingly, only three other genes were significantly upregulated (log-2-fold change >1, adjusted p-value <0.05): *vc0512/aerB, vc0514*, and *vc0515* (**Fig. 4A, Table S3**). We validated these findings using *lacZ* transcriptional reporters. Plasmid-mediated *verA* overexpression was sufficient to induce *P*_*verA*_*-lacZ* in rich LB medium (**Fig. 4B**), consistent with autoregulation. To remove the autoregulatory effect of native VerA from our analysis, we performed additional measurements in a parent strain lacking VSP-II (and thus native *verA*). These data suggest that loss of Zur binding may lead to only a small increase in *verA* transcription, which is further amplified by a VerA-dependent positive feedback loop.

**Figure 4.**
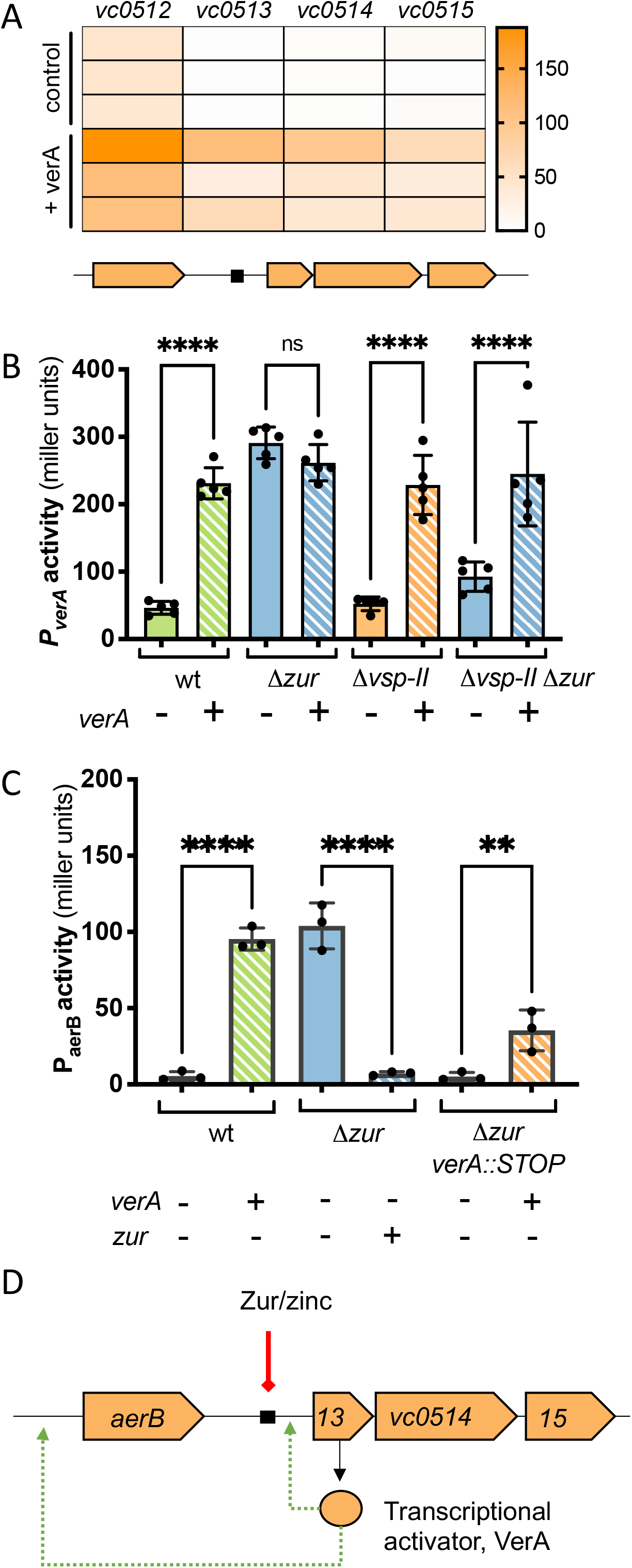
VerA is an AraC-like transcriptional activator that positively regulates *aerB* and the *vc0513-vc0515* operon. (**A**) Overnight cultures of wild-type N16961 carrying either an IPTG-inducible copy of *verA* (+ *verA*) or empty vector (control) were diluted 1:100 in fresh LB containing kanamycin and IPTG (1 mM). RNA was isolated from cells at mid-log phase and prepared for RNA-seq (see Methods and Materials). Heat map shows normalized expression values for differentially expressed genes across three biological replicates. (**B-C**) Overnight cultures strains carrying *lacZ* transcriptional reporters were diluted 1:100 in LB and grown for 3-hr at 37°C. Kanamycin and inducer IPTG (500 µM) were included in the growth medium for *trans* expression (+) of *verA* or *zur* from an IPTG-inducible promoter. (**B-C**) Promoter activity (in Miller Units) was measured via β-galactosidase assays (See Methods and Materials). (**B**) *P*_*verA*_*-lacZ* activity was measured in wild-type, Δ*zur*, Δ*vsp-II*, and Δ*zur* Δ*vsp-II* strains carrying a plasmid-borne, IPTG-inducible copy of *verA* (*+*, striped bars) or an empty vector control (-, solid bars). (**C**) Activity from a P_*aerB*_*-lacZ* reporter was measured in wild-type, Δ*zur*, and Δ*zur verA*::STOP backgrounds harboring a plasmid-borne, IPTG-inducible copy of *verA* or *zur* (+, striped bars) or empty vector control (-, solid bars). (**D**) Proposed model for Zur repression of the *verA* promoter (solid line, red) via a conserved Zur binding site and subsequent VerA-dependent activation (green dashed arrow) of the *aerB* and *verA* promoters.

Interestingly, the *aerB* promoter lacks a conserved Zur binding site; however, our transcriptomic data suggests that Zur-regulated VerA promotes *aerB* transcription. To verify this, we constructed a *P*_*aerB*_*-lacZ* transcriptional reporter. Our initial attempt using a small (400 bp) promoter fragment did not yield detectable signal under inducing conditions (**Fig. S4B-C**). 5’-RACE mapping of the transcription start indicated that *aerB* is part of a much longer transcript (extending >1 kb upstream of the start codon). Thus, we designed a new reporter construct to include this entire region. *P*_*aerB*_ activity in standard LB medium fell below our threshold for detection (**Fig. 4C, Fig. S4C**), but we found that VerA overexpression was sufficient to activate the *aerB* promoter. *P*_*aerB*_ was strongly induced in a Δ*zur* strain background – consistent with our initial RNA-seq – but only if the strain also carried a native or *trans* copy of *verA*. These data indicate that *aerB* expression is dependent upon VerA-mediated activation. In summary, we found that VerA is a Zur-regulated, transcriptional activator that upregulates four genes (*aerB, vc0513-vc0515*).

### AerB *mediates aerotaxis away from air-liquid interface*

Induction of AerB drives *V. cholerae* to aggregate in minimal medium. AerB is predicted to encode a methyl accepting chemotaxis protein (MCP) that senses concentration gradients of a particular ligand (either an attractant or repellent) and relays that signal via Che proteins that alter flagellar rotation (Ud-Din and Roujeinikova 2017). To determine if AerB indeed functions as a chemotaxis receptor, we first tested whether AerB interacts with the chemotaxis coupling protein, CheW. In a bacterial two hybrid assay, AerB and CheW were each fused with one domain of the adenylate cyclase (AC) protein (T18 or T25) and co-transformed into an *E. coli* strain. If the proteins of interest interact, the proximal AC domains will synthesize cAMP and induce *lacZ* expression a cAMP-CAP promoter; thus, a positive protein interaction will yield blue colonies in the presence of X-gal. *E. coli* co-transformed with T18-AerB and CheW-T25 (or the reciprocal tags) yielded bright blue spots (**Fig. 5B)**. We additionally detected strong protein-protein interaction between T18-AerB and T25-AerB, indicating that our chemoreceptor can dimerize (or oligomerize) like other MCPs (Ringgaard et al. 2018). To confirm that AerB’s MCP signaling domain is required for aggregation, we next mutated a glycine residue within the highly conserved C-terminal hairpin loop (R-A-**G**-E/D-X-G) (Alexander and Zhulin 2007) of AerB (**Fig. S5B**), which is required for *in vitro* signal generation in other MCPs (Matthew D Coleman et al. 2005). A Δ*vsp-II* strain expressing AerB[G385C] was unable to aggregate (**Fig. 5A**), consistent with MCP function. Together, these data indicate that AerB indeed functions as a chemotaxis receptor.

**Figure 5.**
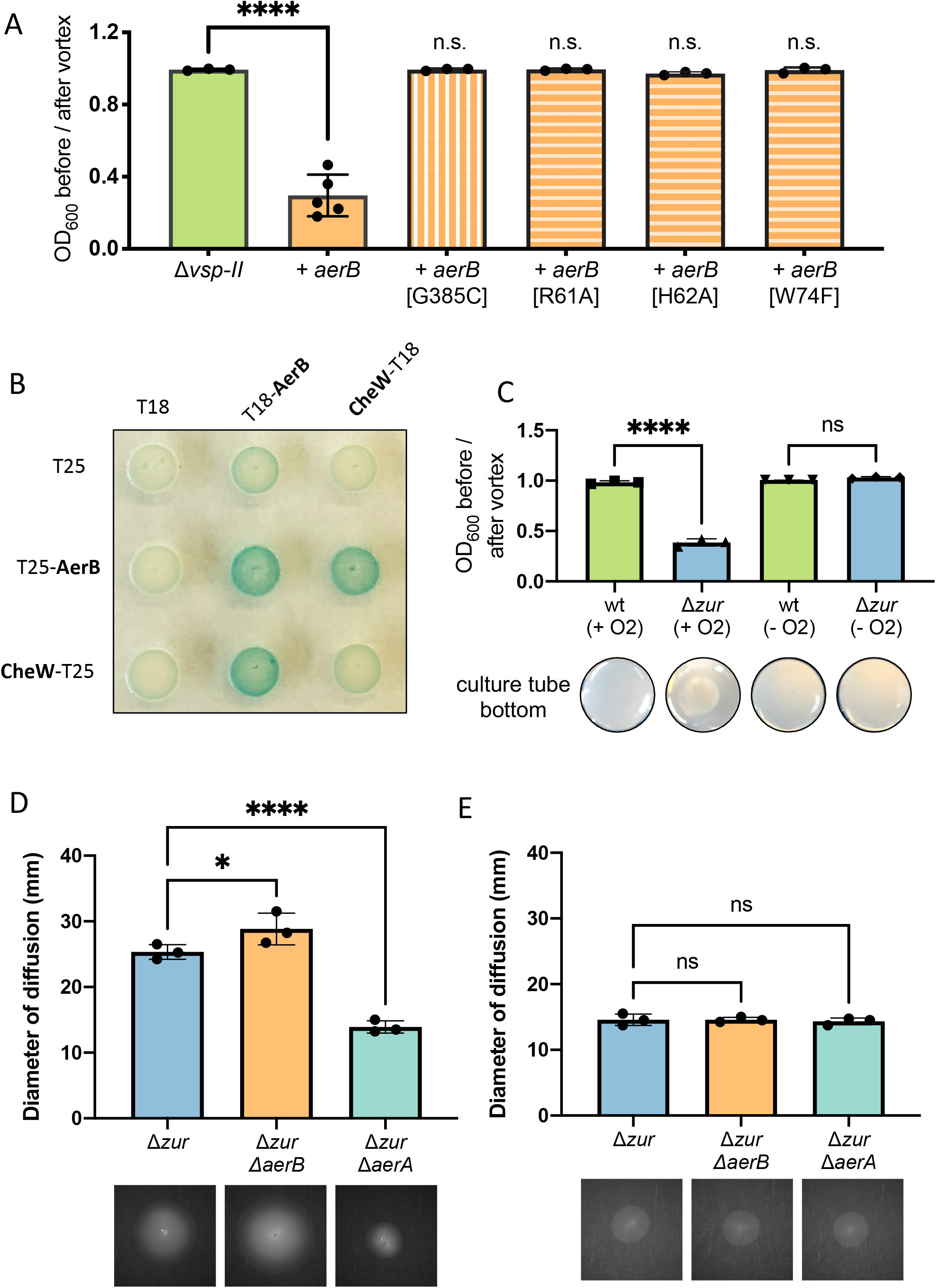
AerB encodes a methyl-accepting chemotaxis protein involved in aerotaxis. (**A**) Δ*vsp-II* strains carrying an integrated, IPTG-inducible copy of either *aerB* or *aerB* point mutants (G385C, R61A, H62A, or W74F) were grown shaking overnight in M9 minimal medium supplemented with glucose (0.2%) and inducer (IPTG, 10 µM) at 30°C. Aggregation was quantified by measuring the optical density (at 600 nm) of the culture supernatant before and after a brief vortex. (**B**) In a bacterial two-hybrid assay, *E. coli* BTH101 was co-transformed with vectors carrying one domain of adenylate cyclase (T18 or T25) or an adenylate cyclase fusion with a protein of interest: CheW-T(18/25) or T(18/25)-AerB. Co-transformants were spotted onto an LB agar containing kanamycin and ampicillin (for selection), X-gal (for blue-white detection), and inducer (IPTG, 500 mM). Plates were incubated overnight at 30°C and for an additional day at room temperature. Blue color signifies positive protein-protein interactions. (**C**) Wild-type and Δ*zur* were grown overnight in 5 mL M9 minimal medium plus glucose (0.5%) plus a terminal electron acceptor (fumarate, 50 mM) and cultured aerobically (+ O_2_). Another set up tubes was prepared anoxically by purging with N_2_ (-O_2_) (see Methods for details). Tubes were grown shaking overnight at 30^°^C and aggregation was quantified as described above. Representative images of the bottom of each culture tube showing pellet presence (for Δ*zur* +O_2_) or absence (all others) are shown. (**D-E**) Strains were grown overnight in LB medium and washed thrice in M9 minimal medium lacking a carbon source. A sterile toothpick was used to inoculate cells into M9 soft agar (0.3%) containing either (**D**) succinate (30 mM) or (**E**) maltose (0.1 mM) as a carbon source. The diameter of diffusion (mm) was measured following a 48-hr incubation at 30°C and representative diffusion patterns are shown for each strain. For all bar graphs, raw data points represent biological replicates, error bars represent standard deviation, and asterisks denote statistical difference via Ordinary one-way ANOVA test (****, p < 0.0001; *, p < 0.05; n.s., not significant).

Intriguingly, the chemical ligands for AerB and the vast majority of *V. cholerae*’s 46 encoded MCPs are yet to be determined (Boin et al. 2004). The AerB N-terminus harbors a PAS domain (Kanehisa et al. 2004), a protein family that typically senses light, oxygen, redox stress, or electron acceptors (Taylor 2007). We hypothesized that the PAS-containing chemoreceptor mediates energy taxis along the oxygen gradient in our vertical culture tubes. We first tested whether oxygen was required for Δ*zur* to aggregate. Wild-type and Δ*zur* were cultured in both aerobic and anoxic tubes in M9 minimal medium with the terminal electron acceptor fumarate to enable anaerobic glucose respiration. Unlike the aerobic cultures, Δ*zur* did not aggregate under anoxic conditions (**Fig. 5C**). A similar result was observed under glucose-fermenting conditions (i.e., when fumarate was omitted from the medium) (**Fig. S6E)**. These data indicate that Δ*zur* aggregation is oxygen-dependent and implicate AerB in energy taxis.

AerB shares 31% amino acid identity with *V. cholerae*’s aerotaxis receptor Aer-2 (renamed here to AerA, as numbers in bacterial gene names can be confused with mutant alleles) (**Fig. S5A**), which exhibits a positive response to oxygen (Boin and Häse 2007). Other homologs include Aer^EC^ (31% identity), which positively responds to oxygen via sensing the electron acceptor FAD (Bibikov et al. 1997; 2000; Taylor 2007). Alignment of AerB with these homologs and Aer^AB^ (from *Azospirillum brasilense* (Xie et al. 2010)) revealed conservation of a critical FAD-binding tryptophan residue, among others (**Fig. S5B**). The corresponding amino acids were mutated in *aerB* and each mutant was expressed in a Δ*vsp-II* background to determine whether they still promoted aggregation. Strains expressing W74F failed to aggregate, suggesting the FAD-binding residue is essential for function (**Fig. 6A**). Two additional mutants (R61A and H62A), corresponding to *E. coli* FAD-binding residues, were also unable to aggregate. The requirement for oxygen and these highly conserved FAD-binding residues suggests that AerB may bind FAD or a similar ligand to facilitate aerotaxis.

**Figure 6.**
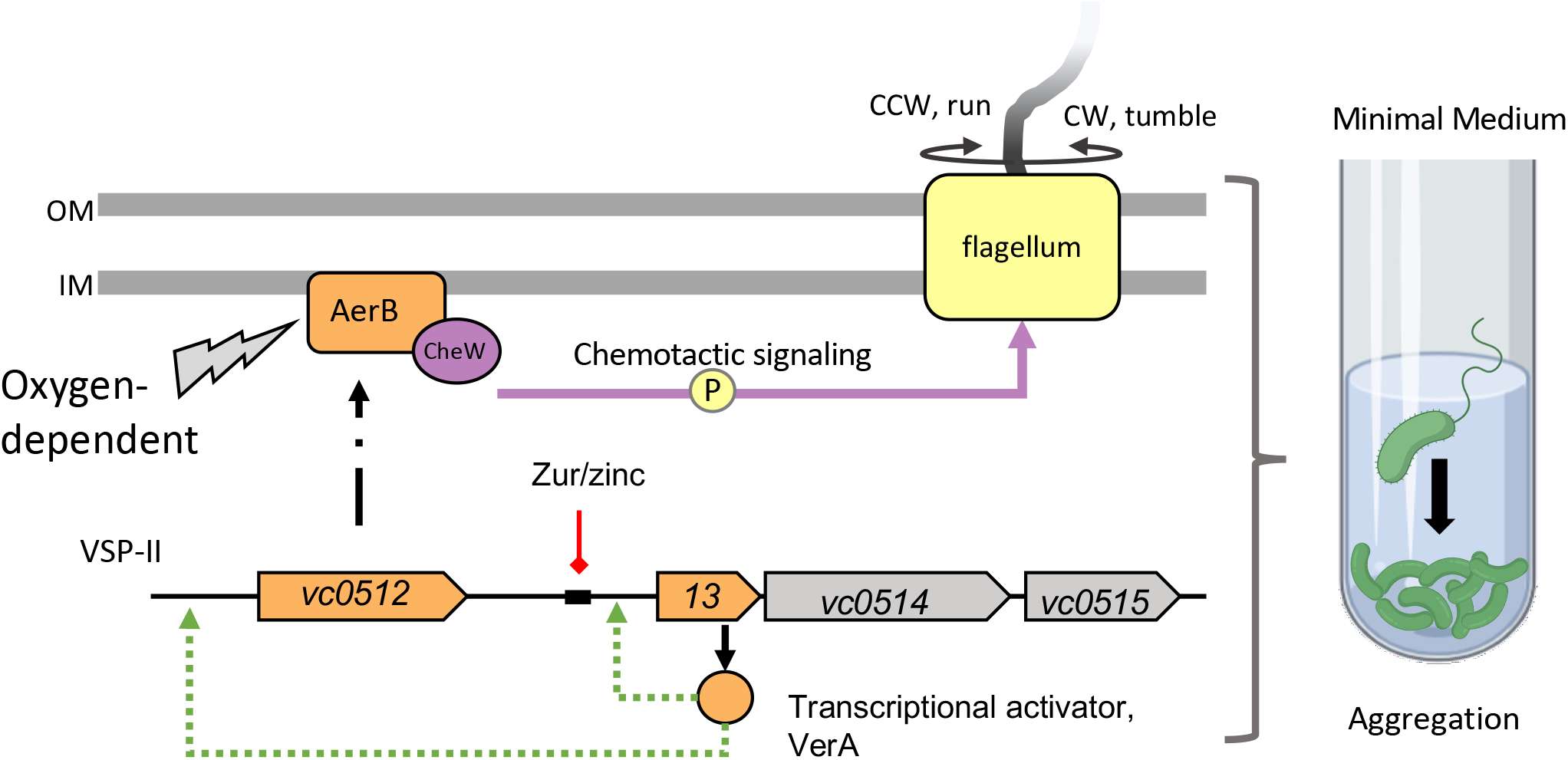
Summary model of Zur-regulation of VSP-II encoded genes the downstream effects on chemotaxis. Zur (orange hexagon) forms a complex with divalent zinc ions (blue circle) and binds with high affinity to specific DNA sequences (black rectangle), repressing transcription of downstream genes (red line). In the absence of *zur* or during zinc starvation, VSP-II genes, including the *vc0513-vc0515* operon are derepressed. The vc0513-encoded transcriptional activator VerA induces transcription (green arrow) of *aerB*, which encodes a chemotaxis receptor. AerB interacts with the chemotaxis coupling protein CheW (purple) and mediates a signal relay that alters flagellar (yellow) rotation and tumbling. AerB induction in minimal medium causes *V. cholerae* to aggregate in an oxygen-dependent manner away from the air-liquid interface.

Based on our aggregation phenotype, we hypothesized that in contrast to AerA and Aer^EC^, AerB appears to mediate a negative response to oxygen and cause cells to accumulate at the bottom of the culture tube. To further interrogate aerotaxis, we examined the swarming dynamics of *V. cholerae* in an established aerotaxis assay, which uses soft agar with carbon sources that vary in their ability to accentuate aerotaxis behavior (Bibikov et al. 1997; Boin and Häse 2007). Succinate, for example, can only be catabolized via respiration, which consumes oxygen, thereby generating an O_2_ gradient that increases with distance from the inoculation site. Since no other classical attractants/repellents are present, motility on succinate plates reveals aerotaxis as the primary taxis behavior (Bibikov et al. 1997). In contrast, maltose agar provides other cues for chemotaxis (including chemotaxis towards maltose itself), obscuring an aerotactic response. The diameter of diffusion in succinate and maltose (at 30°C) was measured two days post-inoculation. All assays were performed in a Δ*zur* background to ensure robust expression of *aerB* from the native promoter and *aerA* mutants were included as a control. Compared to Δ*zur*, the diameter of Δ*zur* Δ*aerB* migration on succinate was significantly increased (**Fig. 5D, Fig. S6A-B**). This is consistent with AerB promoting a negative response to oxygen. Conversely, Δ*zur* Δ*aerA* showed a significant decrease in swarming ability, consistent with AerA promoting a positive response to oxygen, as previously reported (Boin and Häse 2007). (**Fig. 5E, Fig. S6C-D**). In contrast to succinate, there were no significant differences between the swarm diameter of Δ*zur* and the *aer* mutants on maltose swarming plates. These assays were additionally performed in a wild-type background and *aerB* showed no effect on swarming behavior, consistent with lack of *aerB* transcriptional expression in wild-type background (**Fig. S6A-D**). These results corroborate AerB’s function as an aerotaxis receptor and suggest that it mediates an inverse response relative to AerA.

### A model for oxygen-dependent V. cholerae aggregation in zinc starved environments

In summary, we propose the following model for Δ*zur* aggregation in M9 minimal medium (**Fig. 6**). In zinc rich conditions, Zur acts as a repressor of the VerA-encoding *vc0513-vc0515* operon on VSP-II. In the absence of Zur or in zinc starvation, the VerA transcriptional activator induces expression of its own operon and the nearby *aerB*. AerB serves as a receptor for oxygen-dependent energy axis and relays changes in signal concentration to the core chemotaxis machinery and the flagellum. This facilitates aggregation at the bottom of the culture tube in an oxygen-dependent manner.

## Discussion

The mysterious Vibrio Seventh Pandemic Island (VSP-II) present in the El Tor biotype has largely evaded characterization due to lack of knowledge of stimuli that favor its induction. We report that Zur, the transcriptional repressor of the zinc starvation response, is a direct and indirect regulator of numerous VSP-II genes. Novel Zur targets reported here include the *vc0513-vc0515* operon, which encodes the VerA transcriptional activator that increases expression of VSP-II chemotaxis and motility-related genes. One of these secondary targets, AerB, encodes a chemoreceptor involved in aerotaxis and autoaggregation.

### The role of zinc availability in VSP-II induction

It has long been suspected that VPS-II functions as either a pathogenicity or environmental persistence island; however, we and others have not yet identified a set of growth conditions under which VSP-II confers a competitive advantage (O’Shea 2004; Taviani et al. 2010; Sakib et al. 2018; Pant et al. 2020) (**Fig. S7)**. Given the robust expression of VSP-II loci in the absence of Zur, we propose two contexts where *V. cholerae* may encounter zinc starvation and express these island-encoded genes: within the human host, and/or on (chitinous) biotic surfaces in aquatic reservoirs. The human host a well-studied example of a metal-limited environment. Vertebrate hosts sequester desirable metal cofactors (e.g. zinc) in order to restrict the growth of potentially harmful bacteria (i.e. nutritional immunity, (Palmer and Skaar 2016; Kehl-Fie and Skaar 2010; Hood and Skaar 2012; Hennigar and McClung 2016)). Pathogens lacking zinc acquisition systems often exhibit colonization defects *in vivo* (Davis et al. 2009; Sheng et al. 2015; Ammendola et al. 2007; Campoy et al. 2002; Bobrov et al. 2017; Nielubowicz et al. 2010; Sabri et al. 2009; Corbett et al. 2012; Hood et al. 2012), potentially because they are unable to compete against the microbiota for precious metal cofactors (Gielda and DiRita 2012). Induction of zinc starvation genes in pathogenic *V. cholerae* appears to be dependent upon the animal model used; for example, the primary zinc importer and the *vc0513-vc0515* operon are upregulated in a mouse but not in a rabbit model (relative to LB) (Mandlik et al. 2011). Deletion of *V. cholerae*’s zinc importers led to only modest colonization defects in a mouse infection model (Sheng et al. 2015). More generally, *V. cholerae* experiences metal starvation within thick bacterial communities and thus metal transporters and regulators contribute optimal *V. cholerae* biofilm formation (Mueller et al. 2007). Zur-regulated genes (including *vc0503* and *vc0513-vc0515*) are reportedly induced by exposure to chitin oligomers (Meibom et al. 2004), raising the possibility that *V. cholerae* is zinc-limited while colonizing copepods or crustaceans in the environment. It is thus plausible that the Zur-regulated VSP-II genes may be expressed in either of *V. cholerae*’s distinct lifestyles.

### VSP-II-encoded genes facilitate chemotactic responses

Connections between zinc homeostasis and altered motility patterns have been reported in other bacteria, but these phenotypes appear to be indirect consequences of zinc availability rather than Zur repression of secondary transcriptional regulators (Lai et al. 1998; Nielubowicz et al. 2010; Ammendola et al. 2016; Yeo et al. 2017). The VerA-regulated chemoreceptor, AerB, generates aggregation in liquid culture and appears to mediate energy taxis. This is in apparent contradiction with a report that did not find a role for AerB (referred to as *Aer-1*) in aerotaxis (Boin and Häse 2007); however, this may be explained by a lack of native *aerB* expression under their experimental conditions. The balanced action of aerotactic responses conferred by AerA and AerB may function analogously to the Aer and Tsr receptors (Rebbapragada et al. 1997), which enable *E. coli* to navigate to an optimum oxygen concentration.

Although the role of chemotaxis in autoaggregation has not been previously reported in *V. cholerae*, this phenomenon has been characterized in several distantly related bacteria (Alexandre 2015). *A. brasilense*, for example, aggregates in response to oxygen/redox stress via a PAS-containing chemoreceptor homologous to AerB (33% amino acid identity, **Fig. S5A**) (Xie et al. 2010; Bible et al. 2008; Russell et al. 2013). This aerotaxis system may enable *V. cholerae* to avoid redox stress in low-zinc environments, since deletion of zinc importer systems is associated with heightened redox susceptibility in *E. coli* (Sabri et al. 2009). We speculate that AerB may allow *V. cholerae* to colonize other niches within the host (e.g., anaerobic parts of the gut), similar to a redox-repellent chemotaxis system in *Helicobacter pylori* that enables gland colonization *in vivo* (Collins et al. 2016; 2018). Alternatively, this chemotaxis system may allow *V. cholerae* to exploit different niches within the aquatic reservoir (e.g., anoxic sediments with chitin detritus); however, each of these biologically relevant conditions are difficult to recapitulate *in vitro*.

Chemotaxis enhances virulence in a number of enteric pathogens, but this does not seem to generally hold true for *V. cholerae* (Butler and Camilli 2005). In an infection model, non-chemotactic (counter-clockwise biased) mutants outcompeted wild-type *V. cholerae* and aberrantly colonized parts of the upper small intestine (Butler and Camilli 2004), suggesting that chemotaxis is dispensable and possibly deleterious for host pathogenesis. *V. cholerae* appears to broadly downregulate chemotaxis genes in a mouse infection model (Mandlik et al. 2011) and in stool shed from human patients (Merrell et al. 2002). Intriguingly, this decrease may be mediated in part by VSP-I; the island-encoded DncV synthesizes a cyclic AMP-GMP signaling molecule that decreases expression of chemotaxis genes and enhances virulence (Davies et al. 2012). Specific chemoreceptors, however, are upregulated within a host and/or enhance virulence (see (Matilla and Krell 2017) for a review). Given the conflicting roles for chemotaxis within a host, we alternatively suggest that VSP-II encoded chemotaxis genes may serve a purpose in an aquatic environment with oxygen and nutrient gradations.

### Stress and starvation responses can be co-opted by acquired genetic elements

We report that targets of the Zur-regulated VerA appear to be restricted to VSP-II, at least under the conditions tested. This restriction is logical given that transcriptional activators require specific DNA-binding sequences, and these may not be present in the native chromosome of a horizontal transfer recipient. Zur control of secondary regulators, including those that impact gene expression more broadly via signaling molecules (i.e., cyclic di-GMP phosphodiesterases like VC0515) may function to expand the complexity and tunability of the Zur-regulon in response to zinc availability and compounding environmental signals.

Horizontal acquisition of genomic islands can help bacteria (and pathogens) evolve in specific niches. VSP-II retains the ability to excise to a circular intermediate in N16961, indicating the potential for future horizontal transfer events (Murphy and Boyd 2008). We observed that 35% of the ORFs on the prototypical VSP-II island are expressed in the absence of Zur. Intriguingly, genomic island “desilencing” in response to zinc starvation has been reported in diverse bacterium, including *Mycobacterium avium* ssp. Paratuberculosis (Eckelt et al. 2014) and *Cupriavidus metallidurans* (Bütof et al. 2017).

**In summary**, investigation of our Zur-associated aggregation phenotype enabled identification of novel components of the zinc starvation response present on the El Tor Vibrio Seventh Pandemic Island-II. Further characterization of these island-encoded genes may aid in establishing VSP-II’s role as either a pathogenicity or environmental persistence island.

## Methods

### Bacterial growth conditions

Bacterial strains were grown by shaking (200 rpm) in 5 mL of LB medium at 30°C (for *V. cholerae* and *E. coli BTH101*) or 37°C (for other *E. coli*) in borosilicate glass tubes, unless otherwise specified. M9 minimal medium with glucose (0.2%) was prepared with ultrapure Mili-Q water to minimize metal contamination. Antibiotics, where appropriate, were used at the following concentrations: streptomycin, 200□µ□ml^−1^; ampicillin, 100 µg□ml^−1^, and kanamycin, 50 µg□ml^−1^. IPTG was added to induce P_iptg_ promoters at indicated concentrations.

### Plasmid and strain construction

For all cloning procedures, N16961 gDNA was amplified via Q5 DNA polymerase (NEB) with the oligos summarized in **Table S1**. Fragments were Gibson assembled (Gibson et al. 2009) into restriction-digested plasmids. For gene deletions, 700 bp flanking regions were assembled into XbaI-digested pCVD442 (Amp^R^). For complementation experiments, genes of interest were amplified with a strong ribosome binding site and assembled into SmaI-digested pHLmob (kan^R^) and pTD101 (Amp^R^) downstream of an IPTG-inducible promoter. *lacZ* transcriptional reporters were built by amplifying the desired promoter region and assembling into NheI-digested pAM325 (Kan^R^). The resulting promoter-*lacZ* fusions were amplified for assembly into StuI-digested pJL1 (Amp^R^). Cloning for bacterial two-hybrid assays are described in a separate section below. All assemblies were initially transformed into DH5alpha λ*pir* and subsequently into an *E. coli* donor strain (MFD λ*pir* or SM10 λ*pir*). For conjugations into *V. cholerae*, stationary phase recipients and MFD λ*pir* donor strains were washed of antibiotics, mixed in equal ratios, and spotted onto an LB DAP plate. After a 4-h incubation at 37°C, cells were streaked into a selective plate (LB plus ampicillin or kanamycin) to select for transconjugants. Conjugations using SM10 λ*pir* donors were performed in the absence of DAP and with the addition of streptomycin to selective plates. pHLmob transconjugants were purified on an additional kanamycin LB agar plate. Integration vectors were cured through two rounds of purification on salt-free sucrose (10%) agar. Gene deletions or STOP codon replacements introduced by pCVD442 were verified by PCR using the oligos indicated in **Table S1**. Successful integration of *lacZ* targeting vectors (pTD101, pJL1) were identified by blue-white screening on plates containing 5-Bromo-4-chloro-3-indolyl-β-d-galactopyranoside (X-Gal, 40□µg ml^−1^). pJL-1 vectors were additionally checked via PCR. All strains used in this study are summarized in **Table S1**. *V. cholerae* strains were derived from N16961, unless otherwise indicated as E7946 (Miller et al. 1989), C6706 (not strep^R^) (Thelin and Taylor 1996), or Haiti (Son et al. 2011). The N16961 accession numbers for genes referenced in this study are as follows: *zur/vc0378, znuABC/vc2081-vc2083, zrgABC/vc2551-2553, ribA/vc1263, rpmE2/vc0878, rpmJ2/vc0879, shyB/vc0503, aerB/vc0512, verA/vc0513, fliC/vc2199, motB/vc0893, cheA-2/vc2063, cheW-1/vc2059, cheZ/vc2064, cheY/vc2065, vpsL/vc0934, csqS/vca0522, csqA/vc0523, tdh/vca0885, luxS/vc0557, luxQ/vca0736*, and *aerA/vca0658*.

### Site-directed mutagenesis

Site-directed mutagenesis was performed using NEB kit #E0554S according to manufacturer instructions. A pTD101 plasmid carrying *aerB* was used as the template for Q5 amplification with the following mutagenic primer pairs: R61A, SM-1294/1295; and H62A, SM-1296/1297 (**Table S1**). Products were purified and treated with kinase, ligase, and Dpn1 at 37°C for 30 minutes. This reaction mixture was transformed into DH5α □pir. Mutations were confirmed via Sanger Sequencing. *aerB* fragments containing W74F (SM-1306) and G395C (SM-1307) were chemically synthesized by Integrated DNA Technologies (IDT) and assembled into pTD101; sequences are listed in **Table S1**).

### Aggregation assays

Bacterial aggregation was quantified by measuring absorbance (OD600) in a spectrophotometer of the culture supernatant before and after a brief vortex (5 seconds). Aggregation score represents the ratio of before and after pellet disruption; a ratio closer to one indicates that the culture is homogenous, a ratio closer to zero indicates that the cells are concentrated at the bottom of the culture tube.

### Transposon Mutagenesis Screen & Arbitrary PCR

*V. cholerae* N16961 Δ*zur* was mutagenized with Himar1 mariner transposons via an SM10 λ*pir* donor strain carrying pSC189 (Chiang and Rubin 2002). Four independent Δ*zur* transposon libraries were generated, as previously described (Murphy et al. 2019). Each library was separately harvested from the plate using sterile rubber scrapers, vortexed into LB, and preserved in glycerol at −80°C. Individual culture tubes containing 5 mL of M9 minimal medium with glucose (0.2%) and kanamycin were inoculated with transposon libraries. Overnight cultures were back-diluted 1000-fold into fresh medium and incubated overnight; this process was repeated until no visible pellet had formed. Isolated colonies were tested to verify that they did not generate a pellet. The second screen was described identically to the first, except that cultures were first inoculated into a M9 motility plate (0.3% agar) and allowed to migrate for 48 hours at 30°C degrees. Scrapings from the outer zone (collected with a 1 mL pipette tip) were inoculated a culture tube containing M9 minimal medium. For both screens, the transposon insertion site for each isogenic colony was identified by arbitrary PCR (O’Toole et al. 1999). As described elsewhere, this technique amplifies the chromosomal DNA adjacent to the mariner transposon. Amplicons were Sanger sequenced at the Cornell Institute of Biotechnology, and regions of high-quality were aligned to the N16961 reference genome using BLAST (Altschul et al. 1990).

### RNA-seq and analysis

Overnight cultures of wild-type N16961 and the Δ*zur* mutant were diluted 1:100 into LB and grown shaking at 37°C until cells reached mid-log phase (optical density at 600 nm [OD_600_], 0.5). RNA was extracted using mirVana™ miRNA Isolation Kit (Invitrogen, AM1560). Genomic DNA contamination was removed through two DNAfree (Ambion) treatments each followed by glass fiber column purification. Library preparations, Illumina sequencing, and data analysis were performed by GENEWIZ (South Plainfield, NJ). Differentially expressed genes were those with log 2-fold change >1 and an adjusted p-value <0.05.

For VC0513 overexpression, wild-type N16961 carrying either pHLmob or pHLmob(*P*_*iptg*_*-vc0513*) was sub-cultured into LB kanamycin IPTG (500 mM) and grown at 37°C for 3 hour (∼ mid-log phase). Total RNA isolations and DNase treatments were performed as described above. Library preparations, Illumina sequencing, and data analysis (using DESeq2 (Love et al. 2014)) were performed by the Cornell Transcriptional Expression Facility. Differentially expressed genes were those with log 2-fold change >1 and an adjusted p-value <0.05.

### 5’-Rapid Amplification of cDNA Ends (5’-RACE)

Transcription start sites were identified with 5′-RACE. To obtain *vc0512, vc0513*, and *vca1098* transcripts, the Δ*zur* mutant was grown in LB at 37°C until cells reached mid-log phase (optical density at 600 nm [OD_600_], 0.5) and RNA extractions and DNAse treatments were performed as described for RNAseq. PCR was performed to check for genomic DNA contamination; no amplicons were detected within 34 cycles. Reverse transcription was performed with the Template Switching Reverse Transcriptase enzyme mix (NEB #M0466) according to manufacturer protocols using gene specific primers (*vc0512*, SM-1133; *vc0513*, SM-1131; *vca1098*, SM-1129) and the Template Switching Oligo (TSO). PCR Amplification of 5’-transcripts was performed with diluted cDNA, Q5 Hot Start High-Fidelity Master Mix (NEB #M0494), TSO-specific primer, and gene-specific primers (vc0512, SM-1134; vc0513, SM-1132, *vca1098*, SM-1130). Products were sanger sequenced using the following primers: SM-1134, SM-1156, and SM1157 for *vc0512*, SM-1132 for *vc0513*, and SM-1130 for *vca1098*. Primer sequences are listed in **Table S1**.

### β-galactosidase activity measurements

*V. cholerae* strains carrying promoter-*lacZ* fusions were grown overnight in LB at 30°C, with kanamycin for plasmid maintenance. Strains were diluted 1:100 into LB containing Kan and IPTG (1 mM) and grown shaking at 37°C. Exponential phase cells were harvested (∼ 3hr) and β-galactosidase activity against an ortho-Nitrophenyl-β-galactoside substrate (ONPG) substrate was quantified as described elsewhere (Miller 1972; Zhang and Bremer 1995).

### Motility Assays

Motility plates (0.3% agar) were prepared with M9 minimal medium with variable carbon sources (succinate, 30 mM; and maltose, 0.1 mM). Strains were grown overnight in LB medium and washed three times in M9 without a carbon source. Plates were inoculated via toothpick stabs and incubated 30°C 48 hours. The migration diameter (mm) was recorded.

### Bacterial two hybrid assays

Protein-protein interactions were detected using the BACTH bacterial two hybrid system (Karimova et al. 1998). *cheW* and *aerB* (excluding transmembrane domains and native start/stop codons) were cloned into SmaI-digested pUT18(C) (Kan^R^) or pK(N)T25 (Amp^R^) expression vectors to yield N-terminal T(18/25)-Aer or C-terminal CheW-T(18/25) fusions, respectively. Electrocompetent *E. coli* BTH101 were co-transformed with a pUT18 and pKT25 vector that carried either: an unfused adenylate cyclase domain (T18 or T25), the CheW-T(18/25) fusion or the T(18/25)-AerB fusion. Following 1 hr of outgrowth in SOC at 30°C, 10 µL of concentrated outgrowth was spotted onto LB agar containing kanamycin and ampicillin (for selection), X-gal (for blue-white detection), and inducer (IPTG, 500 mM). Plates were incubated overnight at 30°C and for an additional day at room temperature before being imaged.

### Anaerobic Cultures

5 mL of M9 minimal medium without MgSO_4_, CaCl_2_, or carbon source were added to glass culture tubes. Tubes were sealed with rubber stoppers, crimped, purged for 10 cycles (20 sec vacuum, 20 sec N_2_ purge), and autoclaved (gravity, 20 min). Post-autoclaving, the medium was amended with sterile solutions of MgSO_4_ (to 2 mM), CaCl_2_ (to 0.1 mM), glucose (to 0.5%) and with or without fumarate (to 50mM) using sterile syringes and needles. Tubes were injected with a *V. cholerae* cell suspension and grown overnight shaking at 30°C. Aerobic tubes containing M9 glucose (0.5%) with or without fumarate were included as a control. Aggregation was measured via spectrophotometry, as described above.

## Acknowledgements

We thank Sean Murphy and Dr. Daniel Buckley for providing assistance with anoxic culturing, and Dr. John Mekalanos and Dr. Melanie Blokesch for generously sharing *V. cholerae* strains. We thank members of the Dörr lab for helpful discussions. This study was in part supported by NIH grant R01GM130971 to TD and by the Cornell Institute of Host-Microbe Interactions and Disease (CIHMID) (CML, BAJ).

## Figure Legends

**Supplemental Movie 1. A *V. cholerae* N16961** Δ***zur* mutant aggregates at the bottom of culture tubes**. Δ*zur* was grown overnight shaking (200 rpm) in M9 minimal medium with glucose (0.2%) at 30°C.

**Supplemental Table 1. Summary of strains and oligos used in this study**. Strains used in this study are listed with unique identifiers (SGM-#). For *E. coli* donor strains, the primers or gene block used to construct each plasmid are listed in the “Oligos” column. Genetic changes introduced into *V. cholerae* were screened using the method indicated in the “Confirmation” column: either PCR using the indicated primers (SM-#), kanamycin resistance, or blue-white screening on X-gal containing plates.

**Supplemental Table 2. Genes differentially expressed in** Δ***zur* relative to wild-type *V. cholera* N16961**. Transcript abundances in Δ*zur* **r**elative to wild-type were measured using RNA-seq (see Methods for details). Gene ID’s and putative ontology (Kanehisa et al. 2004) are shown for all significant (adjusted p-value < 0.05) differentially expressed (log 2-fold change > 1) genes. Positive values represent up-regulation and negative values represent down-regulation in Δ*zur* relative to the wild-type. Superscripts denote (a) a nearby conserved Zur box, (b) location on VSP-I or (c) location on VSP-II.

**Supplemental Table 3. Genes differentially expressed in *a V. cholerae* strain overexpressing VerA**. Transcript abundances in a strain overexpressing VerA (VC0513) **r**elative to an empty-vector control were measured using RNA-seq (see Methods for details). Gene ID’s and descriptions (Kanehisa et al. 2004) are shown for all significant (adjusted p-value < 0.05) differentially expressed (log 2-fold change > 1) genes.

**Figure S1. Targeted genetic mutations exclude the involvement of a variety of genes in** Δ***zur* aggregation in M9 minimal medium**. All strains were grown overnight in M9 minimal medium plus glucose in **(A**) plastic or (**B-G**) borosilicate glass culture tubes. Aggregation was quantified by measuring the optical density (at 600 nm) of the culture supernatant before and after a brief vortex. The following mutants were tested in a Δ*zur* background: **(B)** other putative Zur-regulatory targets (Δ*vca1098-vca1101* or Δ*vca1090-vca1097*), **(C)** chemotaxis genes (*cheA::STOP*, Δ*cheY*, Δ*cheZ*), **(E)** N16961 *hapR*^repaired^, (**F**) type IV pili (Δ*tcpA*, Δ*mshA*, Δ*pilA*, and Δ*vc0504*), biofilm formation genes (Δ*vspL*), quorum sensing genes (Δ*csqA*, Δ*csqS*, Δ*tdh*, Δ*luxS*, or Δ*luxQ*), (**G**) the Vibrio Seventh Pandemic (VSP) island-I (Δ*vc0175-vc0185*), regions of VSP-II *(“*Δ*VSP-II”*, Δ*vc0491-vc0515;* Δ*vc0490-vc0510*, Δ*vc0511*, Δ*vc0512*, Δ*vc0513 or vc0513::STOP*, Δ*vc0514 or vc0514::STOP*, Δ*vc0515*, or Δ*vc0516*). **(D)** A wild-type, Δ*zur*, a rough mutant (*vc0225::STOP*), and a rough mutant harboring deletions for Δ*fliC*, Δ*motB*, or Δ*vsp-II* were assayed for aggregates. Data points represent biological replicates, error bars represent standard deviation, and asterisks denote statistical difference relative to the wild-type strain via (**A**) unpaired t-test or (**B-G**) Ordinary one-way ANOVA (****, p < 0.0001; ***, p < 0.001;).

**Figure S2. Transposon insertions that prevented** Δ***zur* from aggregating in M9 minimal medium**.

**(A)** Table indicating the number of transposon insertions within motility, chemotaxis, and VSP-II genes for the screens (without pre-selection, v.1; and with pre-selection of motile mutants, v.2) described in **Figure 2. (B)** Approximate location of transposon insertions (triangles) determined by arbitrary PCR (O’Toole et al. 1999) and Sanger sequencing are shown.

**Figure S3. Zur-dependent regulation of the *vca1098* promoter**.

**(A)** Diagram of the *vca1098* promoter region annotated the following features: predicted Zur box, red; predicted −10 region, purple (Solovyev and Salamov 2011); transcription start site, +1 (5’-RACE); predicted ribosome binding site, yellow; proposed start codon, green. Asterisks indicate Zur box nucleotides that were altered in mutant reporters described below. **(B)** *vca1098* promoter-*lacZ* transcriptional reporters (*P*_*vca1098*_*-lacZ*, solid bars) or mutated versions (*P*_*vca1098*_^Zur box* A or B^-*lacZ*, striped bars) were inserted into a wild-type or Δ*zur* background harboring a plasmid-borne, IPTG-inducible copy of *zur* (+) or empty vector control (-). Patterned bars indicate a mutated version of the reporter was used, as described in **A**. Strains were grown overnight in LB and kanamycin, diluted 1:100 in fresh media containing inducer (IPTG, 400 µM), and grown for 3 hours at 37°C. Promoter activity (in Miller Units) was measured via β-galactosidase assays (See Methods and Materials). **(C)** Wild-type and Δ*zur* strains carrying P*vca1098-lacZ* or mutant derivatives were streaked onto M9 minimal medium agar with glucose (0.2%), X-gal, and with or without added zinc (ZnSO_4_, 10 µM). Plates were incubated overnight at 30°C and then for an additional day at room temperature. *vca1098* promoter activity is indicated by a blue colony color.

**Figure S4. Transcription start sites and construction of *lacZ* transcriptional reporters**.

(**A**) Diagram of the *verA* promoter regions annotated the following features: predicted Zur box (red), predicted −10 and −35 regions (purple) (SOFTBERRY), suggested start codon (green). (**B**) Schematics for two attempted *P*_*aerB*_*-lacZ* reporters containing either 400 bp or 1,314bp of the promoter region are shown. (**C**) The *P* _*aerB*_ ^400 bp^*-lacZ and P* _*aerB*_ ^1,314 bp^*-lacZ* reporters in a wild-type or Δ*zur* background were struck onto LB X-gal plates and incubated overnight at 30°C and then for an additional day at room temperature. *P*_*aerB*_ expression is indicated by a blue colony color.

**Figure S5. AerB protein alignment with homologs in *V. cholerae, E. coli* and *A. brasilense***.

**(A)** Results of protein BLAST (Altschul et al. 1990) and **(B)** Clustal Omega alignment (Larkin et al. 2007) of AerB against homologs from *V. cholerae* (AerA, encoded by *vca0685*) *E. coli* (Aer^EC^, encoded by *b3072*) and *A. brasiliensis* (AerC^AB^, encoded by *AKM58_23950*). Conserved ligand binding and MCP residues targeted for mutation are indicated by orange and pink arrows, respectively.

**Figure S6. *V. cholerae* swarm assays with chemoreceptor mutants**.

(**A-D**) The indicated strains were grown overnight in LB medium and washed thrice in M9 minimal medium lacking a carbon source. A sterile toothpick was used to inoculate cells into M9 soft agar (0.3%) with either (**A-B**) succinate (30 mM) or (**C-D**) maltose (0.1 mM) as a carbon source. The diameter of diffusion (mm) was measured following a 48-h incubation at 30°C and representative swarms are shown (**A**,**C**). Note: Data for Δ*zur*, Δ*zur* Δ*aerB*, Δ*zur* Δ*aerA, and* Δ*zur* Δ*aerA* Δ*aerB* are the same as shown in **Fig. 5** and are shown here for comparison with a wild-type background. (**E**) Wild-type and Δzur were grown overnight in 5 mL M9 minimal medium plus glucose (0.5%) fermentatively (without a terminal electron acceptor) and cultured under aerobic (+ O_2_) or anoxic (-O_2_) conditions (see Methods for details). Tubes were grown shaking overnight at 30°C and aggregation was quantified via spectrophotometry as described previously. All data points represent biological replicates, error bars represent standard deviation, and asterisks denote statistical difference via Ordinary one-way ANOVA test (****, p < 0.0001; ***, p < 0.001, *, p < 0.05, and n.s., not significant).

**Figure S7. Presence of VSP islands does not impact growth in zinc-chelated medium**. Wild-type, Δ*vsp-I*, and Δ*vsp-II* were grown overnight in M9 minimal medium with glucose (0.2%) at 30°C. Cultures were washed twice and diluted 1:100 into (A) fresh M9 minimal medium glucose (0.2%), (**B**) plus the zinc-specific chelator TPEN (250 nM), or (**C**) plus TPEN and exogenous zinc (ZnSO_4_, 1 µM). Growth at 30°C of each 200-µl culture in a 100-well plate was monitored by optical density at 600 nm (OD_600_) on a Bioscreen C plate reader (Growth Curves America).

